# Mating changes the genital microbiome in both sexes of the common bedbug *Cimex lectularius* across populations

**DOI:** 10.1101/819300

**Authors:** Sara Bellinvia, Paul R. Johnston, Susan Mbedi, Oliver Otti

## Abstract

Many bacteria live on host surfaces, in cells, and specific organ systems. Although gut microbiomes of many organisms are well-documented, the bacterial communities of reproductive organs, i.e. genital microbiomes, have received little attention. During mating, male and female genitalia interact and copulatory wounds can occur, providing an entrance for sexually transmitted microbes. Besides being potentially harmful to the host, invading microbes might interact with resident genital microbes and affect immunity. While sexual transmission of individual symbiont species is relatively well-understood, few studies have addressed how mating changes genital microbiomes. Here, we characterize male and female genital microbiomes in four different populations of the common bedbug *Cimex lectularius* and investigate mating-induced changes. We dissected reproductive organs from virgin and mated individuals to sequence their genital microbiomes. We show that mating changes genital microbiomes, suggesting bacteria are sexually transmitted. This raises the question how the host’s physiological and immunological mechanisms control mating-induced changes in their genital microbiomes. Also, genital microbiomes varied between populations and the sexes. This provides evidence for local and sex-specific adaptation of bacteria and hosts, again suggesting that bacteria might play an important role in shaping the evolution of reproductive traits. Coadaptation of genital microbiomes and reproductive traits might further lead to reproductive isolation between populations, giving reproductive ecology an important role in speciation. Future studies should investigate the transmission dynamics between the sexes and populations to uncover potential reproductive barriers.

## INTRODUCTION

Animals have intimate associations with bacteria. These bacteria are found on host surfaces, within and between host cells, or are associated with specific organ systems, such as the gut^1–3^. Also, reproductive organs of a variety of animals often harbor several different bacterial species. Microbes have been found on male reproductive organs and in semen of vertebrates^4–9^, but also in female reproductive organs of birds and mammals, both including humans^5,10–13^. In insects, besides intracellular reproductive manipulators (reviewed in^14^), extracellular microbes have been found both on the male copulatory organ and within the female reproductive organs (reviewed in^15^, also see^16,17^). Interestingly, the microbiomes of whole body homogenates or the gut are sex-specific in a variety of species^18–20^. Possible reasons could be differences in behaviors or feeding strategies, different functions of the two sexes in the ecosystem, or different roles in reproduction. Supporting the latter, pronounced differences exist between the genital microbiomes of female and male red-winged blackbirds^5^ and bedbugs^16,17^, respectively. However, the role of genital microbes in reproduction and the exact composition of these genital microbiomes is still unknown for most insect species.

Microbiomes are dynamic and react to environmental change. Diet^21,22^, age^23^, climate^24^, and even time of day^25^ alter the composition of the gut microbiome of humans, mice and lizards. Some life-history events affect multiple microbiomes at the same time, for instance pregnancy changes the vaginal, gut, and oral microbiomes^26^. Genital microbiomes might be subject to similar changes. For instance, women have a distinct vaginal microbiome before and after the menopause^27^. An obvious pattern that has the potential to affect genital microbiomes is mating as the microbiomes of both sexes encounter each other and potentially interact. Therefore, physiological changes in anticipation to the production of offspring might occur. To date, little is known about the potential effects of mating on the composition of genital microbiomes, especially in species other than humans.

Microbes are sexually transmitted in a large range of species (e.g.^28–30^). Not only typical sexually transmitted microbes (STM) are transferred during mating, also opportunistic microbes (OM) inhabiting the genitalia^17,31,32^ might be transferred. Possible entry ports are genital openings and copulatory wounds, which frequently occur for example in insects, spiders, flatworms and gastropods^33^. These OM might have a more direct effect compared to STM: Once transferred into the female genital tract, sperm will be exposed to a rich microbial flora^7,10^. Several bacterial species, such as *Escherichia coli*^34–36^, *Pseudomonas aeruginosa*^37^, and *Staphylococcus aureus*^38^ can decrease sperm motility and agglutinate spermatozoa in humans. In insects, bacteria from the environment have the potential to increase sperm mortality^39^. Therefore, male reproductive success might depend on the microbe communities within the female, and males would be selected to protect their sperm. Females could be invaded by STM or sexually transmitted OM, disturbing the present genital microbiome. The intestinal microbiota of mice, for instance, was shown to be altered by *Salmonella typhimurium*^40^ and *Citrobacter rodentium*^41^. Copulatory wounding during mating further increases the risk of opportunistic infection not only in invertebrate species^42,43^, but also in humans^44^. Both males and females might be subject to a considerable cost of reproduction, even if OM do not always become pathogenic. Hosts should invest in reducing or regulating the number of STMs or sexually transmitted OM to prevent uncontrolled growth, most likely by using their immune system. However, sexually transmitted bacteria disturbing the composition of the genital microbiome might select not only for a host immune response, but also for defensive responses in the resident microbiota^16^. Therefore, the host and its endosymbionts have a mutual interest in keeping invading bacteria in check. In some organisms, the resident microbiota is part of the interaction with invading microbes^45–48^.

If genital microbiomes are subject to OM, an adaptation to environmental microbes seems conceivable. Indeed, the human vaginal microbiome seems to depend on ancestry: American women of African, Asian, European, and Hispanic ancestry harbored distinct vaginal microbiomes^11^ and African ancestry was related to larger alpha diversity of the vaginal microbiome compared to European ancestry^49^, raising the question whether there are conserved differences in the community composition between populations. Despite the potential relevance for speciation, not much is known about the adaptation of insect genital microbiomes to different host populations which might lead to reproductive isolation. Another unresolved question is how genital microbiomes are affected by mating and whether induced changes differ between populations. Unlike in humans such mating-induced changes can be experimentally investigated in insects with relatively little effort^17^.

Bedbugs are an interesting system to study mating-induced changes of insect genital microbiomes because several organs are involved in reproduction. During mating, these organs might be invaded by microbes that use the routes of ejaculate transfer as entrance to the reproductive organs. The male ejaculate consists of spermatozoa that are stored in the male sperm vesicles and seminal fluid from the seminal fluid vesicles. After mounting the female, the male transfers the ejaculate via its copulatory organ, the paramere, into the female paragenital copulatory organ, the mesospermalege^50^. This female immune organ is filled with immune cells (hemocytes) and has evolved to reduce the cost of mating associated with bacterial infection^51^. After a few hours, sperm travel through the hemolymph towards the ovaries^50^. All tissues involved could be invaded by microbes that impose a risk of infection or sperm damage. We predict female tissues to be more affected by an invasion of OM compared to male tissues because bacteria could be transmitted via the paramere and enter the immune organ via copulatory wounds. We expect the reproductive organs to differ in their microbial composition and the microbial composition itself to change due to mating. Indeed, there are potential indications of sexual transmission in bedbugs^17^. Furthermore, microbial communities differ between reproductive organs and copulation increases the similarity of female and male organs. Bacteria present in mated but not virgin individuals of one sex are found in the opposite sex and resident bacteria are replaced with introduced bacteria^17^. Our aim is to investigate whether these findings are a general pattern across bedbug populations.

Here, we describe the genital microbiome of male and female common bedbugs *Cimex lectularius*. We focus on a potential difference in the microbial community between populations, between the two sexes, between organs, and between virgin and mated individuals to evaluate the effect of the external environment on the genital microbiome and the risk of a change in composition via copulation. We expect distinct microbiomes between the investigated groups. If compositional changes due to mating are caused by sexual transmission, we expect to find bacterial strains that occur in virgin individuals of only one sex but in mated individuals of both sexes. Mating-induced changes in the genital microbiome should be caused by a loss or introduction of bacterial strains or by a replacement of resident with invading bacterial strains. We predict that the mechanisms differ between females and males due to the different function of the organs involved in mating. In case members of a specific bacterial strain invade the genital microbiome, mating should change the abundance of the given bacterial strain. To test our predictions, we use a community ecology approach based on 16S rRNA sequencing data from the genital microbiomes of four bedbug populations.

## RESULTS AND DISCUSSION

We sequenced 643 samples from bedbug reproductive organs or cuticle to characterize the composition of the genital microbiomes and investigate the effect of mating. After filtering, we obtained 149 sequence variants (SVs) from 495 samples (Table 1). Average alpha diversity given by the Simpson index (1-D) was 0.59 (0.56; 0.62) (mean and 95% confidence interval) (Figure S1).

**Table 1.** Sample sizes and number of bacterial communities that were successfully sequenced for each group of samples. Sampled organs were: cuticle (C), sperm vesicle (S), seminal fluid vesicle (Sf), paramere (P), mesospermalege (M), hemolymph (H) and ovary (O).

| Population | Sex | Organ | Mating status | N total | N sequenced |
| --- | --- | --- | --- | --- | --- |
| A | Male | C | Virgin | 10 | 7 |
| A | Male | C | Mated | 10 | 6 |
| A | Male | S | Virgin | 10 | 7 |
| A | Male | S | Mated | 11 | 7 |
| A | Male | Sf | Virgin | 10 | 7 |
| A | Male | Sf | Mated | 10 | 6 |
| A | Male | P | Virgin | 10 | 7 |
| A | Male | P | Mated | 10 | 6 |
| A | Female | C | Virgin | 11 | 7 |
| A | Female | C | Mated | 11 | 8 |
| A | Female | M | Virgin | 10 | 8 |
| A | Female | M | Mated | 10 | 6 |
| A | Female | H | Virgin | 11 | 8 |
| A | Female | H | Mated | 10 | 6 |
| A | Female | O | Virgin | 10 | 6 |
| A | Female | O | Mated | 9 | 7 |
| B | Male | C | Virgin | 10 | 8 |
| B | Male | C | Mated | 10 | 8 |
| B | Male | S | Virgin | 10 | 8 |
| B | Male | S | Mated | 10 | 10 |
| B | Male | Sf | Virgin | 10 | 5 |
| B | Male | Sf | Mated | 10 | 7 |
| B | Male | P | Virgin | 10 | 5 |
| B | Male | P | Mated | 10 | 7 |
| B | Female | C | Virgin | 10 | 8 |
| B | Female | C | Mated | 10 | 10 |
| B | Female | M | Virgin | 10 | 9 |
| B | Female | M | Mated | 10 | 7 |
| B | Female | H | Virgin | 10 | 9 |
| B | Female | H | Mated | 10 | 10 |
| B | Female | O | Virgin | 10 | 9 |
| B | Female | O | Mated | 10 | 9 |
| C | Male | C | Virgin | 10 | 9 |
| C | Male | C | Mated | 10 | 8 |
| C | Male | S | Virgin | 10 | 10 |
| C | Male | S | Mated | 10 | 9 |
| C | Male | Sf | Virgin | 10 | 6 |
| C | Male | Sf | Mated | 10 | 10 |
| C | Male | P | Virgin | 12 | 7 |
| C | Male | P | Mated | 8 | 6 |
| C | Female | C | Virgin | 10 | 10 |
| C | Female | C | Mated | 10 | 8 |
| C | Female | M | Virgin | 10 | 8 |
| C | Female | M | Mated | 10 | 7 |
| C | Female | H | Virgin | 10 | 6 |
| C | Female | H | Mated | 10 | 7 |
| C | Female | O | Virgin | 10 | 6 |
| C | Female | O | Mated | 10 | 8 |
| D | Male | C | Virgin | 10 | 9 |
| D | Male | C | Mated | 10 | 7 |
| D | Male | S | Virgin | 10 | 9 |
| D | Male | S | Mated | 10 | 9 |
| D | Male | Sf | Virgin | 10 | 7 |
| D | Male | Sf | Mated | 10 | 8 |
| D | Male | P | Virgin | 10 | 7 |
| D | Male | P | Mated | 10 | 9 |
| D | Female | C | Virgin | 10 | 9 |
| D | Female | C | Mated | 10 | 8 |
| D | Female | M | Virgin | 10 | 8 |
| D | Female | M | Mated | 10 | 8 |
| D | Female | H | Virgin | 10 | 9 |
| D | Female | H | Mated | 10 | 9 |
| D | Female | O | Virgin | 10 | 8 |
| D | Female | O | Mated | 10 | 8 |

### Microbiomes of virgin bedbugs

#### Differences in the composition of cuticular and genital microbiomes

Virgin bedbugs harbored distinct cuticular and genital microbiomes. The members of microbial communities and their relative abundance differed between the cuticle and the internal reproductive organs (F_1,218_=5.257, R^2^=0.024, q=0.003) but not between the cuticle and the external reproductive organ, the paramere (F_1,91_=1.411, R^2^=0.015, q=0.14) or the internal reproductive organs of both sexes and the paramere (F_1,177_=2.350, R^2^=0.013, q=0.02) (Figure S2). The groups did not differ in between-individual difference (F_2,243_=1.521, p=0.22).

On the first glance, the compositional differences between microbiomes of internal reproductive organs and cuticle seem to be caused by location. Environmental bacteria have been found on the external reproductive organ of bedbugs^32^, suggesting cuticle and paramere should mostly be colonized by environmental bacteria. In contrast, internal organs might be more protected from environmental bacteria but at the same time they might harbor symbionts that play a role in bedbug reproduction. Although we found distinct microbial communities on the cuticle and in the internal reproductive organs, the microbiome of the paramere was not different from the microbiome of all other organs. Hence, its microbiome could be influenced by the microbiomes of both the cuticle and the internal microbiome. Alternatively, a combination of organ location and function might be responsible for this finding.

#### Composition of genital microbiomes

Genital microbiomes of virgin bedbugs harbored on average 25 (21; 28) SVs (mean and 95% confidence interval) and were specific to each group of bedbugs. The members of microbial communities and their relative abundances differed between populations (F_3,170_=1.885, R^2^=0.030, p=0.004) and sexes (F_1,170_=3.776, R^2^=0.020, p=0.001) (Figure S3). Even different organs taken from the same sex harbored distinct microbiomes (F_4,170_=1.578, R^2^=0.034, p=0.008). Between-individual variation did not differ between populations (F_3,175_=1.410, p=0.24), sexes (F_1,177_=2.138, p=0.15), or organs (F_5,173_=1.488, p=0.20).

We did not find any SV present in all samples of a given sex, but three SVs occurred in at least half of all individuals of both sexes. Males and females did not differ in the prevalence of a gammaproteobacterial endosymbiont of *C. lectularius* (males: 67%, females: 64%) and one *Rickettsia* strain (males: 58%, females: 59%), whereas more females than males harbored a second *Rickettsia* strain (males: 59%, females: 69%). The relative abundance of the three most prevalent genera varied tremendously from 0% to 50% (gammaproteobacterial endosymbiont) or from 0% to 100% (both *Rickettsia* strains) in individual samples. Virgin males and females shared 69% (population A), 88% (population B), 86% (population C), and 95% (population D) of all SVs found in virgin bedbugs from the specific population (Figure 1).

**Figure 1.**
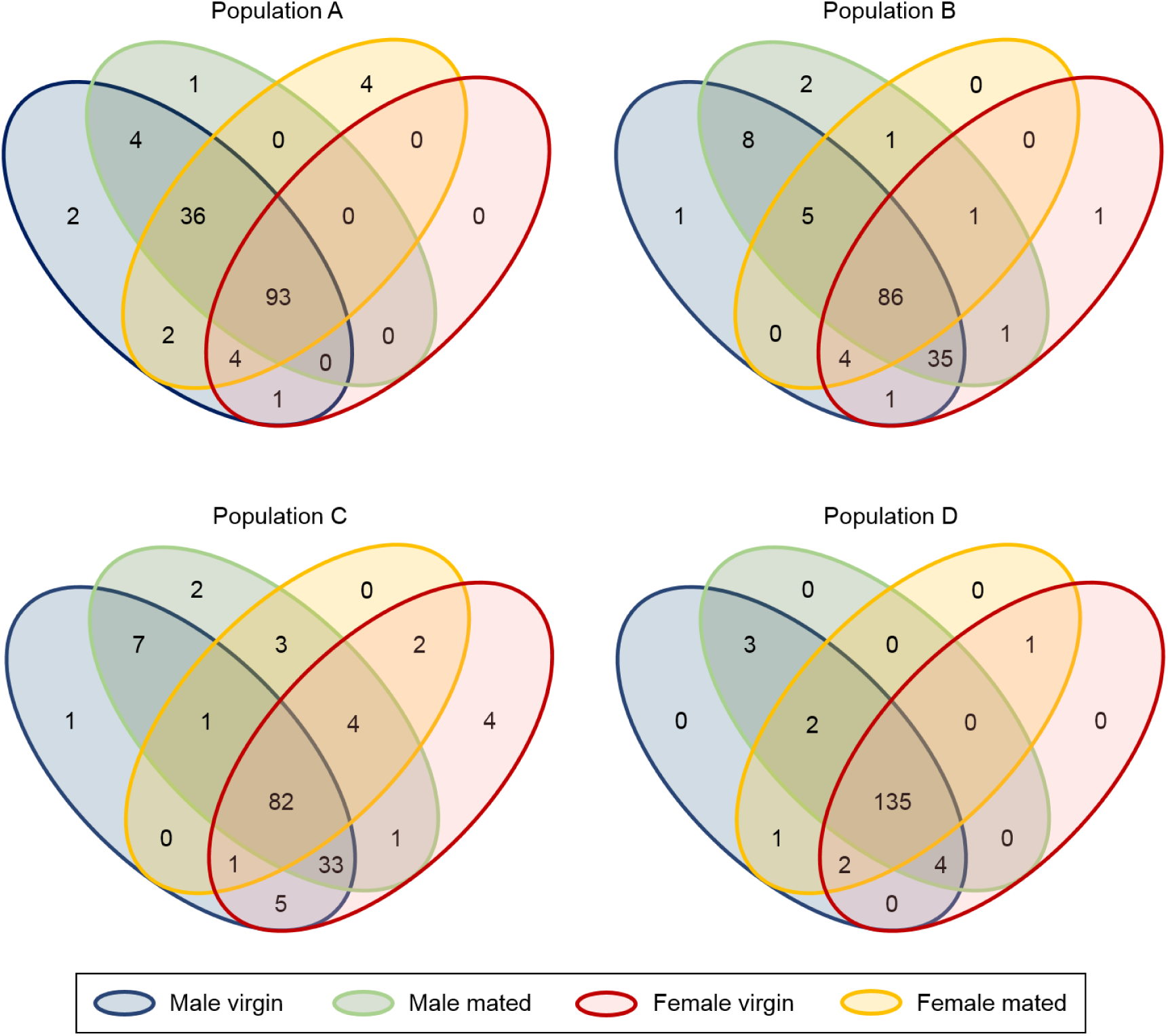
Occurrence of all SVs within the genital microbiomes split up by population, sex, and mating status. A total of 149 SVs were detected, of which 147 were harbored by population A, 146 by population B, 146 by population C, and 148 by population D.

Population-specific microbiomes are likely to arise if host and microbes coevolve. Sexual conflict and genetic drift might play a role as well. Ultimately, differences in microbiome composition might be involved in speciation processes such as reproductive isolation. We here present a system under controlled lab conditions, fed with the same food source, nevertheless being held in separate containers. Despite of the controlled environmental factors, we found differences in microbiome composition between bedbug populations. Our findings are in accordance with the evidence for ancestry-dependent vaginal microbiomes in humans^11,49^. In contrast to our study under controlled lab conditions, it is difficult to exclude lifestyle factors, such as diet, sex partners, etc. as the cause of differences in vaginal microbiomes. We are aware that this comparison with humans might be far-fetched, as the taxa clearly differ in their interaction with microbes. Unfortunately, little is known about genital microbiomes in insects. We could draw here parallels to gut microbiomes because they are also heavily influenced by influx of environmental microbes, but this is equally hypothetical. Therefore, we prefer to highlight that our study shows that insects and other arthropods are ideal study systems to experimentally investigate genital microbiomes and their role in reproduction in more detail.

Differences in microbiome composition between females and males have been found in whole body homogenates or gut samples in a variety of species^18–20^. These differences could be explained by different behaviors or feeding strategies, different functions of the two sexes in the ecosystem, or different roles in reproduction. Despite a lack of studies investigating the origin of sexual dimorphism in the microbiome, pronounced differences exist between the genital microbiomes of female and male red-winged blackbirds^5^ and bedbugs^16,17^, respectively. Furthermore, our finding of organ-specific bacterial communities within the same sex is in accordance with the varying microbiome composition along the female reproductive tract in humans^52^. Either the different accessibility for OM or the function of the organs could be responsible for such a specificity. To our knowledge, we provide evidence for organ-specific genital microbiomes in female insects for the first time.

### Mating-induced changes in the genital microbiomes

#### Changes in the presence and relative abundance of specific SVs

We found mating-induced changes in the genital microbiomes of females and males regarding the presence of specific SVs and their relative abundance as virgin and mated individuals harbored distinct genital microbiomes (F_1,355_=1.938, R^2^=0.005, p=0.03) (Figure S3). We found no interaction of organ and mating status (F_5,317_=1.003, R^2^=0.013, p=0.49), population and mating status (F_3,317_=0.762, R^2^=0.006, p=0.83), or population, organ and mating status (F_15,317_=0.779, R^2^=0.031, p=0.99). Between-individual variation was similar between mating status (F_1,363_=0.010, p=0.92), organs (F_5,359_=1.462, p=0.20), and populations (F_3,361_=0.281, p=0.84).

In all populations, males and mated females shared bacterial strains that were not present in virgin females (Figure 1). In two out of four populations, mated males harbored bacteria that were present in females but not in virgin males. This does not necessarily indicate sexual transmission of these strains since their prevalence did not differ between virgin and mated individuals of the same sex (q>0.05). All bacteria found in mated but not virgin individuals that were not present in the reproductive organs of the opposite sex occurred on the cuticle of virgin bedbugs.

Changes in strain composition can be caused by direct sexual transmission of OM, via ejaculate transfer or being transferred from the male paramere or the female cuticle into the mesospermalege. Genital openings or copulatory wounds provide both entry and exit for bacteria. Indeed, our results indicate that bacteria that occur in mated but not in virgin individuals of one sex originate from the reproductive organs of the opposite sex as well as from the cuticle. However, we found no significant differences in the prevalence of these strains between virgin and mated individuals. A classical transmission study could show whether sexual transmission and transmission of OM from the environment is a common phenomenon in bedbugs.

#### Potential mechanisms affecting composition

We partitioned beta diversity, given as the Jaccard index, into nestedness (strain loss or introduction of new strains) and turnover (replacement of strains) to analyze the potential mechanism of the mating-induced change found in genital microbiomes. In 16 out of 24 cases the proportion of beta diversity explained by SV turnover was higher than the proportion explained by SV nestedness (Table 2). In male organs, turnover was the dominant mechanism in 10 out of 12 cases, while in females both mechanisms accounted for the composition change in half of the cases.

**Table 2.**
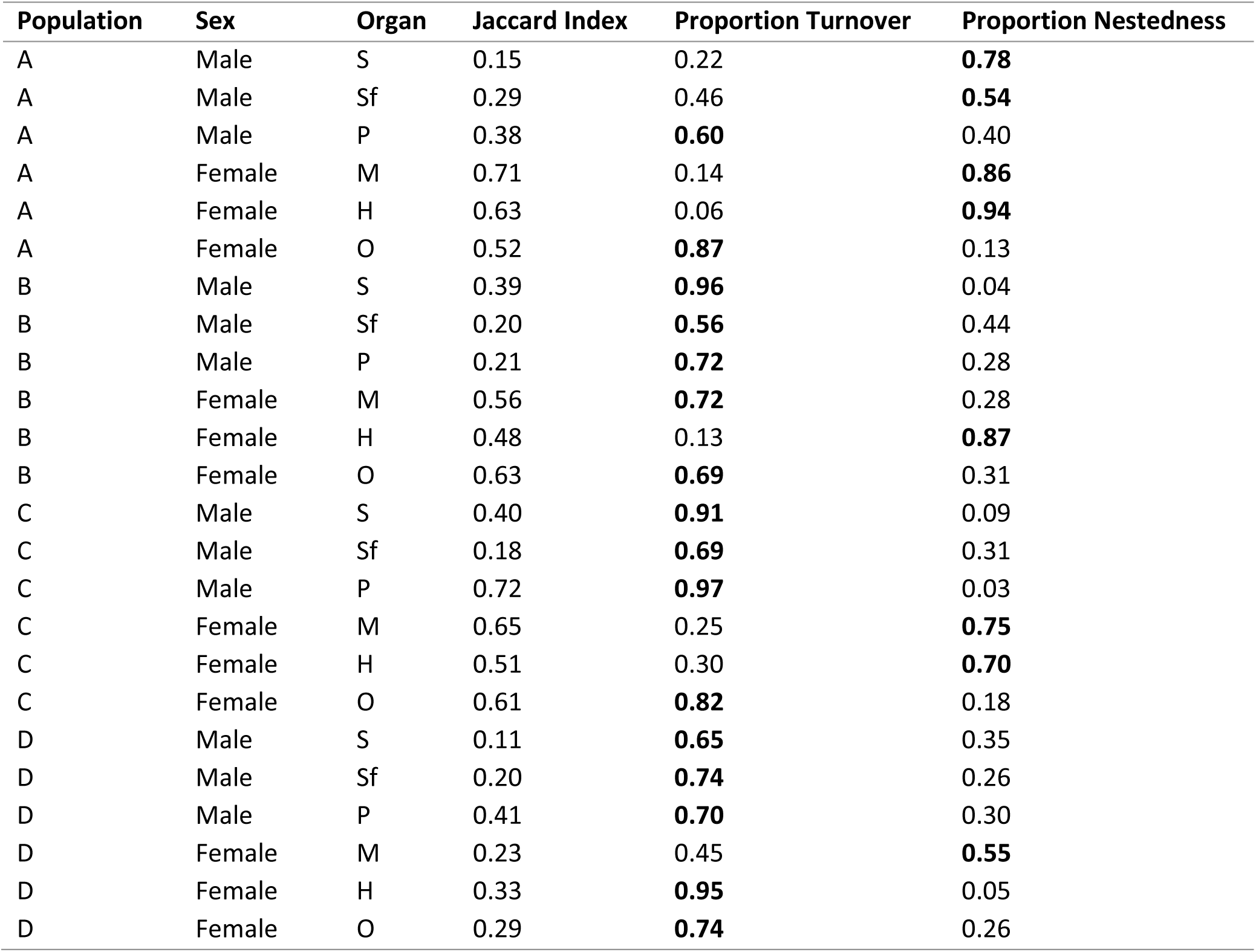
Turnover (replacement of strains) and nestedness (strain loss or introduction of new strains) of the microbiomes of the sperm vesicle (S), the seminal fluid vesicle (Sf), the paramere (P), the mesospermalege (M), the hemolymph (H) and the ovary (O) obtained by partitioning beta diversity (Jaccard index) with the *vegan* package^69^. The mechanisms explaining a higher proportion of beta diversity within population and organ are indicated in bold type.

| Population | Sex | Organ | Jaccard Index | Proportion Turnover | Proportion Nestedness |
| --- | --- | --- | --- | --- | --- |
| A | Male | S | 0.15 | 0.22 | **0.78** |
| A | Male | Sf | 0.29 | 0.46 | **0.54** |
| A | Male | P | 0.38 | **0.60** | 0.40 |
| A | Female | M | 0.71 | 0.14 | **0.86** |
| A | Female | H | 0.63 | 0.06 | **0.94** |
| A | Female | O | 0.52 | **0.87** | 0.13 |
| B | Male | S | 0.39 | **0.96** | 0.04 |
| B | Male | Sf | 0.20 | **0.56** | 0.44 |
| B | Male | P | 0.21 | **0.72** | 0.28 |
| B | Female | M | 0.56 | **0.72** | 0.28 |
| B | Female | H | 0.48 | 0.13 | **0.87** |
| B | Female | O | 0.63 | **0.69** | 0.31 |
| C | Male | S | 0.40 | **0.91** | 0.09 |
| C | Male | Sf | 0.18 | **0.69** | 0.31 |
| C | Male | P | 0.72 | **0.97** | 0.03 |
| C | Female | M | 0.65 | 0.25 | **0.75** |
| C | Female | H | 0.51 | 0.30 | **0.70** |
| C | Female | O | 0.61 | **0.82** | 0.18 |
| D | Male | S | 0.11 | **0.65** | 0.35 |
| D | Male | Sf | 0.20 | **0.74** | 0.26 |
| D | Male | P | 0.41 | **0.70** | 0.30 |
| D | Female | M | 0.23 | 0.45 | **0.55** |
| D | Female | H | 0.33 | **0.95** | 0.05 |
| D | Female | O | 0.29 | **0.74** | 0.26 |

Against our expectations, there was potential transmission to males, indicated by a replacement of resident with new bacterial strains. This is surprising because it is less likely for the male to be subject to copulatory wounding and the distance between the environment and the internal reproductive organs is larger compared to females. Thus, we would have expected invading microbes to be flushed out via ejaculate transfer rather than to invade the internal reproductive organs. However, in case males do not apply pressure to transfer the ejaculate, bacteria might reach the internal organs through the ejaculatory duct via a capillary effect similar to the invasion of human testicles by urethral pathogens through the seminal duct or the epididymal duct^53^. The induced expression of antimicrobial peptides in the genital tract of *Drosophila* males in response to bacteria deposited on the genital plate^54^ suggests bacteria can enter the reproductive organs of male insects via genital openings. Experimental manipulation of bacteria on the paramere might clarify whether and how bacteria can enter the paramere and move through the ejaculatory duct towards the sperm vesicles and seminal fluid vesicles.

#### Abundance changes

Mating changed the abundance of several bacterial strains in almost all organs within populations (Table 3). However, the proportion of differentially abundant SVs did not differ between populations (F_3,19_=0.421, p=0.74) or between males and females (F_1,19_=1.554, p=0.23) and population did not interact with sex (F_3,16_=1.181, p=0.35). No clear direction of abundance change, i.e. decrease or increase in abundance, for populations, sexes, or organs was identified (Figure 2). The absolute log2-fold change representing the strength of abundance change differed between populations (F_3,316_=5.577, p<0.001) but not between sexes (F_1,316_=0.048, p=0.826) and population did not interact with sex (F_3,313_=0.946, p=0.419).

**Table 3.**
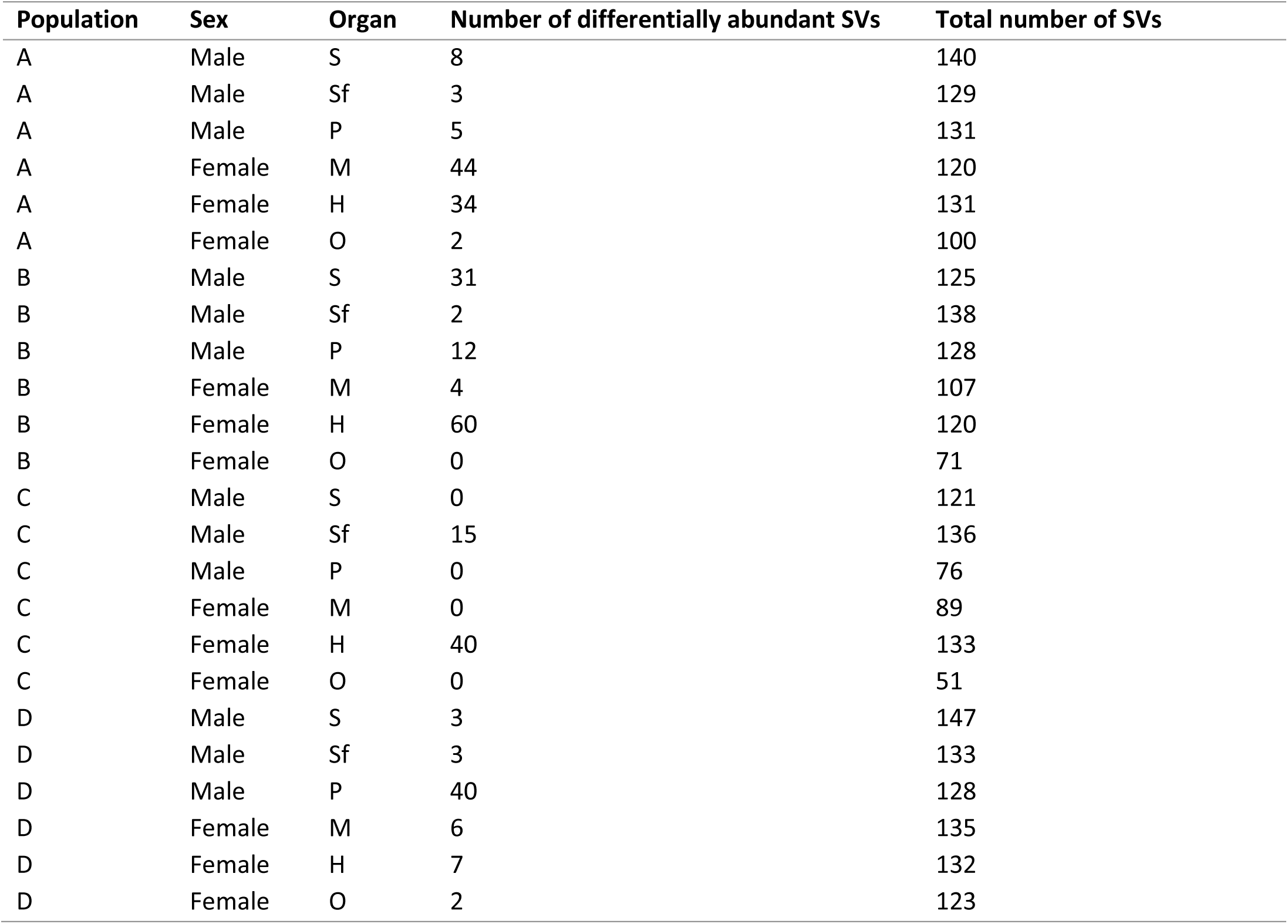
Differential abundance of the microbiomes of the sperm vesicle (S), the seminal fluid vesicle (Sf), the paramere (P), the mesospermalege (M), the hemolymph (H) and the ovary (O) from virgin vs mated individuals. Values were obtained by fitting a GLM in the *edgeR* package^70,75^. Only SVs with significant differential abundance (q<0.05) are reported.

**Figure 2.**
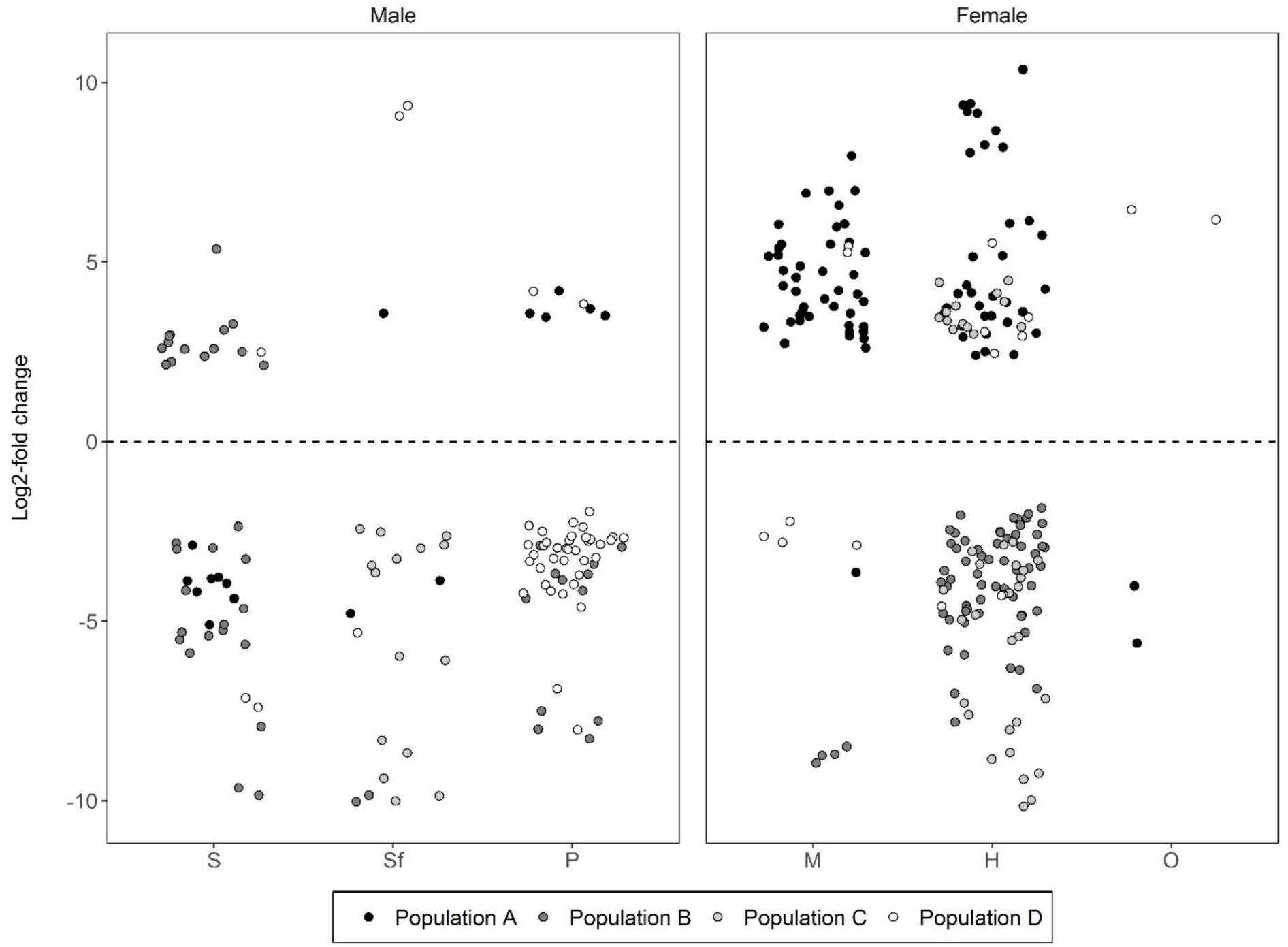
Abundance change of SVs present in the microbiomes of the sperm vesicle (S), the seminal fluid vesicle (Sf), the paramere (P), the mesospermalege (M), the hemolymph (H) and the ovary (O) from virgin vs mated individuals as estimated by GLM fits in the *edgeR* package^70,75^. Given is the log2-fold change of SVs with significant differential abundance as a measure of the direction and strength of abundance change.

We expected that the genital microbiomes of females would be more affected by invading bacteria, but the abundance change or its strength did not depend on sex. A high stability of the genital microbiome is especially important for females because bacteria can enter the body more easily compared to males. Bacteria might not only decrease survival^51^ but also cause a trade-off between immunity and mating and hence decrease fecundity. Such a reduction in fecundity after an immune challenge has been found in dipterans, coleopterans, and orthopterans^55^ and is likely to also exist in bedbugs. Moreover, bacteria harm sperm within the female directly. *Escherichia coli*, *Pseudomonas aeruginosa*, and *Staphylococcus aureus* decrease sperm motility^34–38^ and incapacitate spermatozoa^38^ in humans, at least *in vitro*, and environmental bacteria increase sperm mortality in bedbugs^39^. To reduce the costs of mating-associated infections, bedbugs have evolved the mesospermalege^51^. The high number of hemocytes^50^ able to phagocytose bacteria^56^ in this organ might stabilize its microbiome and protect against invading bacteria. In addition, lysozyme in the seminal fluid of males^57^ and in the mesospermalege produced in anticipation of mating^58^ could help to reduce invading bacteria. Furthermore, endosymbionts have been shown to interact with invading microbes^45–48^ and might help to control non-resident bacteria in the genital microbiomes. Future studies should investigate the effect of the species with the largest abundance changes in the genital microbiomes on fecundity and survival and what adaptations have evolved to eliminate the possible threat to host integrity.

We have demonstrated that genital microbiomes of the common bedbug *Cimex lectularius* differ between populations, sexes and organs. Our findings show that genital microbiomes are sensitive to an activity that every sexually reproducing animal has to face, i.e. mating. Future studies should investigate sexual transmission dynamics of OM in combination with fitness effects on both sexes. Experimental manipulation of the female immune system could provide important information about the importance of immunity in response to genitalia-associated bacteria. Finally, coadaptation of genital microbiomes and reproductive traits might lead to reproductive isolation between populations, giving reproductive ecology an important role in speciation.

## METHODS

### Bedbug culture

Common bedbugs (*Cimex lectularius* L.) are an established model system for examining consequences of sexual selection and reproductive physiology^51,59^. We used four large stock populations of the common bedbug. Three populations were field caught from London, UK in 2006 (A), from Nairobi, Kenya, in 2008 (C), and from Watamu, Kenya, in 2010 (D). The fourth population (B) was a long-term lab stock originally obtained from the London School of Hygiene and Tropical Medicine over 20 years ago. The populations were maintained in a climate chamber at 26±1°C, at 70% relative humidity and a light cycle of 12L:12D and fed weekly using the protocol of Hase^60^. After eclosion, all individuals were kept in sex-specific groups of 20-30 individuals in 60ml plastic pots containing filter paper. Males were fed twice. Females were fed three times, with the last feeding on the day of dissection because fully fed females cannot resist copulation via traumatic insemination ^61^.

### Mating and sample preparation

Dissections and DNA extractions were conducted in the laboratory of the Animal Population Ecology at the University Bayreuth. We dissected 643 three-week-old males and females, originating from four different populations (Population A: N=163, population B: N=160, population C: N=160, population D: N=160). Half of the individuals were randomly mated for 60 seconds and dissected 1-2 hours after mating. That means that the sperm were still present in the mesospermalege of mated females at the time of dissection. We used standard dissection techniques minimizing the potential of contamination by a lab butane burner (Labogaz 206, Campingaz, Hattersheim, Germany) placed next to the dissection microscope. The dissection kit was autoclaved each day and each individual forceps and surgical scissors were dipped in ethanol (70%) and flame sterilized before each dissection.

We collected different reproductive tissues and cuticle samples from females and males (for sample sizes see Table 1). Each sample was taken from a different bedbug since it is difficult to obtain all tissues from the same individual. From males we collected sperm vesicles, seminal fluid vesicles and paramere. In females we investigated the mesospermalege, ovaries and hemolymph. Hemolymph was collected using a sterilized glass capillary pulled to a fine point. Each tissue was transferred into an Eppendorf tube containing 150µl of phosphate-buffered saline. We investigated whether invading bacteria in the genital microbiomes of mated bedbugs originated from the paramere or the cuticle by transferring whole females or males, whose paramere had been removed, into a tube. To evaluate contamination during the dissection process, we placed an open Eppendorf tube containing only phosphate-buffered saline next to the dissection microscope while processing the bedbugs. This tube was then processed in the same way as all tissue samples from which we extracted DNA. All samples were frozen in liquid nitrogen and stored at −80°C.

### Molecular methods

Prior to transferring the samples to the MicroBead tubes from the MO BIO UltraClean Microbial DNA Isolation Kit (dianova GmbH, Hamburg, Germany), we homogenized them using pestles made from sterile pipette tips (200µl). We followed the UltraClean protocol, except that we dissolved the DNA in 30µl elution buffer for higher yield. DNA was stored at −20°C. To control for contamination during DNA extraction we performed one extraction without adding any tissue.

The library preparation and sequencing was done in the laboratory of the Berlin Center for Genomics in Biodiversity Research (BeGenDiv). The samples were split up into four sequencing runs, each balanced in terms of population, sex, organ, and mating status, resulting in four libraries with 128±25 (mean±SD) samples each, including controls. Using the universal primers 515fB (5’-GTGYCAGCMGCCGCGGTAA-3’, ^62^) and 806rB (5’-GGACTACNVGGGTWTCTAAT-3’, ^63^), we amplified the variable V4 region from the bacterial 16S rRNA (denaturation: 94°C for 3 min; 30 cycles of denaturation: 94°C for 45 s, annealing: 50°C for 1 min, extension: 72°C for 90 s; extension: 72°C for 10 min). After amplicons had been purified using Agencourt AMPure XP beads (Beckmann Coulter GmbH, Krefeld, Germany), a unique combination of two eight nucleotide long index sequences per sample were used for barcoding each sample in a second PCR (denaturation: 95°C for 2 min; 8 cycles of denaturation: 95°C for 20 s, annealing: 52°C for 30 s, extension: 72°C for 30s; extension: 72°C for 3 min). After a second purification step with AMPure beads, the DNA concentration of the PCR products was quantified with the Quant-iT PicoGreen dsDNA Assay Kit (Invitrogen, Carlsbad, CA, USA). Samples that had a higher concentration than 5 ng/µl were diluted to this concentration, all other samples were left undiluted. The quality of the pooled amplicons was verified with a microgel electrophoresis system (Agilent 2100 Bioanalyzer, Agilent Technologies, Santa Clara, CA, USA). The resulting libraries were sequenced by an Illumina MiSeq sequencer. For each of the two PCRs per sequencing run we had two negative controls containing only purified water instead of DNA. Two additional samples for each PCR per sequencing run contained the bacterial DNA from a whole homogenized bedbug to increase sequencing depth. The sequences were deposited in NCBI’s Sequence Read Archive with the accession number PRJNA560165.

### Bioinformatical analysis

#### Data processing and check for contamination

All data processing and analyses were performed in R (version 5.3.1^64^) with the packages *dada2* (version 1.10.1^65^), *decontam* (version 1.2.0^66^), *phyloseq* (version 1.19.1^67^), *pairwiseAdonis* (version version 0.0.1^68^), *vegan* (version 2.4-5^69^), and *edgeR* (version 3.22.1^70^). We used the *dada2*^65^ pipeline to filter and trim the sequences with the *fastqPairedFilter* function. The first 10 bp were removed and the sequences were truncated after 260 bp (forward reads) or 200 bp (reverse reads), or at the first instance of a quality score less than or equal to 2. Sequences with expected errors higher than 2 were discarded. The remaining sequences were dereplicated with the default parameters of the *derepFastq* function and denoised with the *dada* function (enabled *selfConsist* option). We constructed a sequence variant (SV) table with the *makeSequenceTable* function. All SVs were checked for chimeras with the *removeBimeraDenovo* function and default parameters. The taxonomy was assigned with the Greengenes database^71^. To control for contamination, we scored the controls for dissection, DNA isolation, target PCR and indexing PCR with the *decontam* package^66^ and removed the SVs that were found to be contaminants based on their prevalence in controls. We further removed all SVs that occurred in only one sample and that had less than 0.01% of the total number of reads, as suggested by Caporaso et al.^72^. We verified the taxonomical assignment with Blast2Go^73^ and its default options. We excluded uncultured or environmental samples. If Blast2Go did not find a match, we used NCBI’s BLASTn with the default options, excluding uncultured and environmental sample sequences. The taxonomic assignments of the different algorithms were in accordance for kingdom, phylum, class, order, family and genus level in 110 out of 149 SVs. In one of the mismatches, the BLAST hit with the highest e-value and coverage belonged to an endosymbiont of *C. lectularius*, the unclassified gammaproteobacterium mentioned by Hosokawa *et al.*^74^. We therefore changed the taxonomy assignment of this SV. In seven out of the mis-assigned cases, all BLAST hits agreed on one genus and we therefore changed the assignment. In 38 other cases there was no clear BLAST result. Hence, we kept the Greengenes assignment for the levels that were congruent with the BLAST results and changed the assignment of the other levels to “Unclassified”. Information regarding sample type, read numbers for samples and SVs, and assigned taxa can be found in the Supplement (Table S1-S3).

#### Statistical analysis of differences in the microbiomes of virgin bedbugs between populations, sexes, and organs

We calculated the relative abundance of all genera by dividing the number of reads for the given SV within one sample by the total number of reads for all SVs within the same sample. Alpha diversity was estimated based on the Simpson index (1-D) with the *phyloseq* package^67^. We then analyzed the differences in microbiome composition between internal reproductive organs, external reproductive organs and cuticle with a multilevel pairwise comparison using PERMANOVAs (*pairwiseAdonis* package^68^) based on Bray-Curtis dissimilarities. P-values were adjusted with the Benjamini-Hochberg procedure. Between-individual differences between the three groups were analyzed with an ANOVA of the function *betadisper* in the *vegan* package^69^.

To compare the genital microbiome composition between populations and sexes, we used a PERMANOVA (999 permutations, *vegan* package^69^) with the fixed effects population, sex, and organ nested within sex based on Bray-Curtis dissimilarities obtained with the *phyloseq* package^67^. Between-individual variation was then compared with 3 separate ANOVAs of the *betadisper* function in the *vegan* package^69^ for populations, sexes, and organs. The prevalence of all SVs in males and females was calculated by dividing the number of samples for the given sex that contained a given SV by the total number of samples of the given sex.

#### Statistical analysis of effects of mating on the bacterial communities

Differences in the composition of genital microbiomes from virgin and mated bedbugs were analysed with a PERMANOVA (999 permutations, *vegan* package^69^) with the fixed effects population, organ, and mating status, and their interactions based on Bray-Curtis dissimilarities obtained with the *phyloseq* package^67^. Non-significant interactions were removed from the model. We then compared between-individual variation with separate ANOVAs of the *betadisper* function in the *vegan* package^69^ for population, organ and mating status.

To analyse the potential of sexual transmission, we extracted the SVs present in virgin individuals of only one sex but in mated individuals of both sexes. We applied Fisher’s exact tests to determine whether the prevalence of each of these SVs differed between virgin and mated individuals. P-values were adjusted with the Benjamini-Hochberg procedure.

We analyzed whether the different bacterial strains hosted by virgin and mated individuals are due to a loss or introduction of specific strains (nestedness) or due to a replacement of resident bacterial strains with introduced bacteria (turnover). To analyze the contribution of these two mechanisms, we partitioned beta diversity, in this case given by the Jaccard index, into nestedness and turnover with the function *nestedbetasor* in the *vegan* package^69^. Before applying this function, we produced a presence-absence matrix that included all SVs present in each population, organ, and mating status. With the package *edgeR*^70,75^ we tested for differential abundance of SVs between the genital microbiomes of virgin and mated bedbugs with the *glmQLFit* and *glmQLFTest* function. Normalization factors were calculated with the relative log expression^76^ and applied to raw read counts. Contrasts were built for every mating status within organ and population and p-values were adjusted with the Benjamini-Hochberg procedure and a false discovery rate of 1%. The resulting proportion of SVs with differential abundance was analyzed with a logistic regression with quasibinomial error structure and the fixed effects population and sex, including their interaction. Groups were then compared with an ANOVA. To analyze the effects of population and sex on the strength of abundance change given by the absolute log2-fold change, we fitted a GLM with Gaussian error structure with the fixed effects sex and organ nested within sex and population as the random factor. Normality of the data was restored with the Box-Cox transformation (*MASS* package^77^) and checked visually before groups were compared with an ANOVA. Non-significant interactions were removed from the models.

## ACKNOWLEDGEMENTS

S.B. was supported by a grant of O.O. awarded by the German Research Foundation (OT 521/2-1). We would like to thank M. Kaltenpoth for helpful comments on the manuscript.

## AUTHOR CONTRIBUTIONS

O.O., P.R.J., and S.B. conceived the idea and designed the experiment. S.B. and S.M. carried out the experiment. S.B. and P.R.J. performed the bioinformatics and statistical analysis. S.B., P.R.J., S.M. and O.O. interpreted the results and S.B. and O.O. wrote the manuscript. All authors read and approved of the final manuscript.

## CONFLICT OF INTEREST

The authors have declared no conflict of interest.

## SUPPLEMENTARY INFORMATION

**Figure S1.**
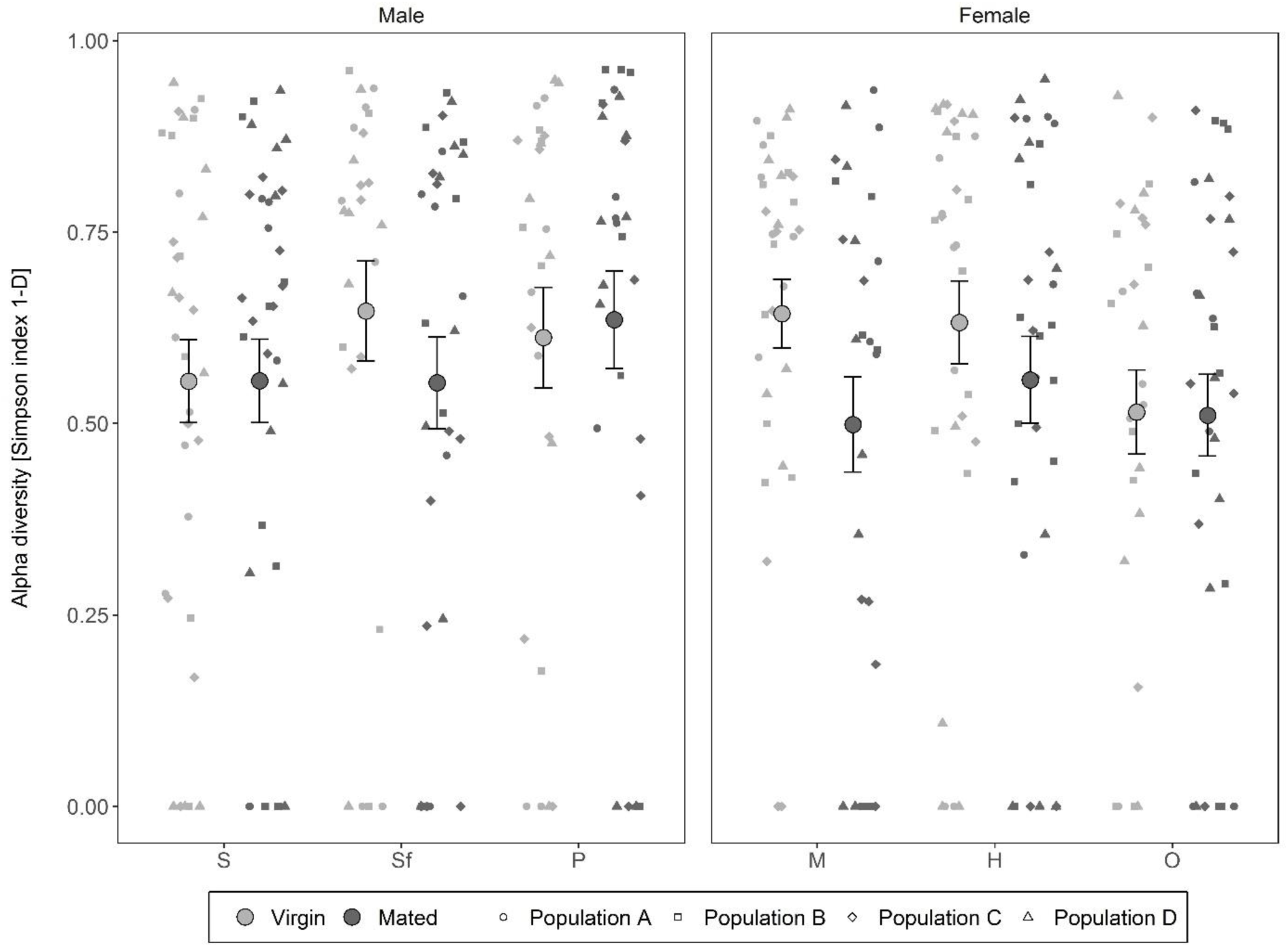
Alpha diversity of each sample in the sperm vesicle (S), the seminal fluid vesicle (Sf), on the paramere (P), in the mesospermalege (M), the hemolymph (H) and the ovary (O). Depicted are means and standard errors of the mean. Simpson indices were obtained with the *estimate_richness* function in the *phyloseq* package^67^.

**Figure S2.**
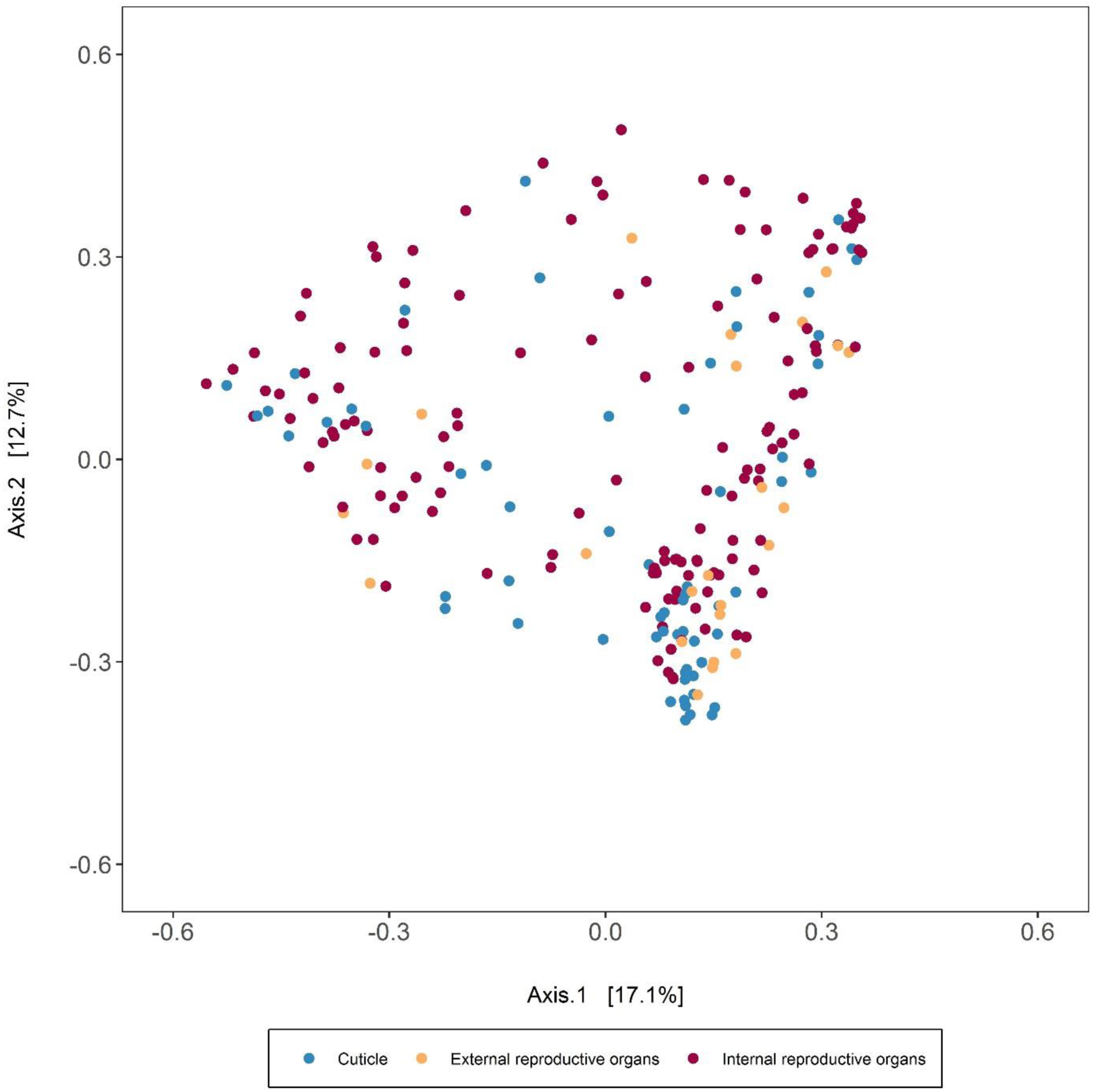
PCoA of microbiomes from cuticle in comparison to the external reproductive organ of males) and internal reproductive organs of both sexes. Ordination was performed with the *ordinate* function and plotted with the *plot_ordination* function in the *phyloseq* package^67^.

**Figure S3.**
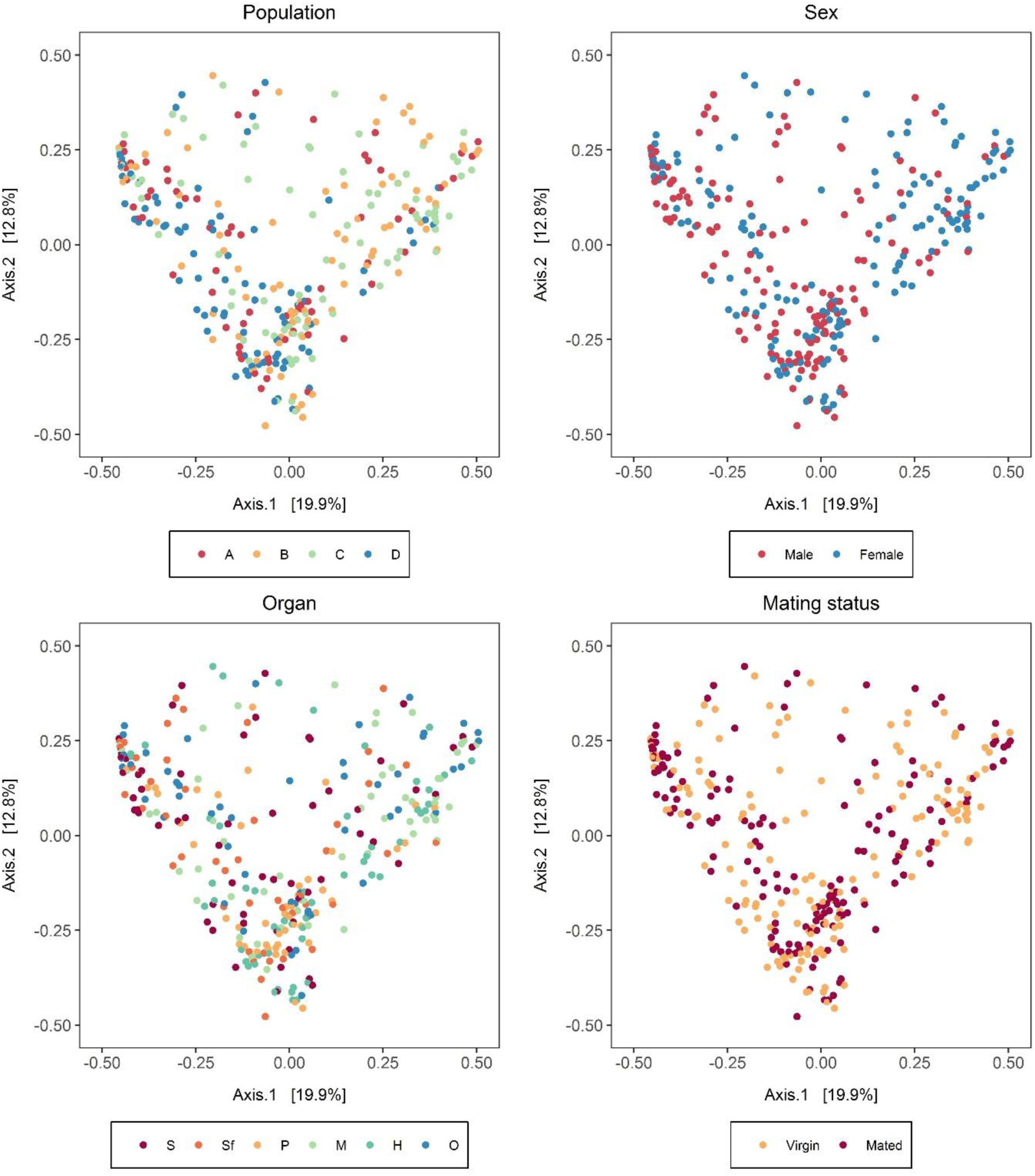
PCoA of all groups of genital microbiomes. Organs are sperm vesicle (S), seminal fluid vesicle (Sf), paramere (P), mesospermalege (M), hemolymph (H) and ovary (O). Ordination was performed with the *ordinate* function and plotted with the *plot_ordination* function in the *phyloseq* package^67^.

All supplemental tables can be requested via email.

**Table S1** Sample information regarding population, sex, organ, mating status, and dissection date.

**Table S2** Raw read counts for each sample and SV.

**Table S3** Assigned taxa for each SV.

